# Dominance Analysis: A formalism to uncover dominant energetic contributions to biomolecular condensate formation in multicomponent systems

**DOI:** 10.1101/2023.06.12.544666

**Authors:** Daoyuan Qian, Hannes Ausserwoger, Tomas Sneideris, Mina Farag, Rohit V. Pappu, Tuomas P. J. Knowles

## Abstract

Phase separation in aqueous solutions of macromolecules is thought to underlie the generation of biomolecular condensates in cells. Condensates are membraneless bodies, representing dense, macromolecule-rich phases that coexist with the dilute, macromolecule-deficient phase. In cells, condensates comprise hundreds of different macromolecular and small molecule solutes. Do all components contribute equally or very differently to the driving forces for phase separation? Currently, we lack a coherent formalism to answer this question, a gap we remedy in this work through the introduction of a formalism we term energy dominance analysis. This approach rests on model-free analysis of shapes of the dilute arms of phase boundaries, slopes of tie lines, and changes to dilute phase concentrations in response to perturbations of concentrations of different solutes. We present the formalism that underlies dominance analysis, and establish its accuracy and flexibility by deploying it to analyse phase spaces probed *in silico, in vitro*, and *in cellulo*.

---

Phase separation in aqueous solutions of proteins and nucleic acids is thought to be a major driver of spatial organisation in cells that gives rise to mesoscale membraneless bodies known as biomolecular condensates [1, 2]. In the simplest scenario, a binary mixture comprising a protein or nucleic acid in a complex aqueous solvent separates into two coexisting phases. Phase separation occurs above a threshold macromolecular concentration to minimise the overall free energy of the system [3]. The concentrations of macromolecules and all components of the complex solvent in the coexisting phases are prescribed by the conditions of phase equilibrium, viz., the equalisation of species-specific chemical potentials and osmotic pressure across phases [3, 4].

In most theories of phase separation and approaches used to analyse experimental data, the contributions of solvent components, including salt, pH, buffer, and crowders, are folded into effective two and three-body interaction parameters [5–10]. Such analysis ignores the possibility of differential inter-phase partitioning of solvent components. We illustrate this point using the work of Bremer et al. [11, 12]. They mapped temperature-dependent phase diagrams for over thirty different variants of A1-LCD, a prion-like low complexity domain from the protein hnRNPA1 [11]. We shall consider two sequences they studied, which they designated as WT (for wild-type) and −12F+12Y. In the latter, all Phe residues (F) were replaced with Tyr (Y). Everything else about the two sequences is identical. At a given temperature and for identical solution conditions, Bremer et al. quantified the threshold concentrations, designated as *c*_sat_, above which each of the protein solutions undergo phase separation. They found that the *c*_sat_ value of −12F+12Y is lower than that of the WT sequence. The inference is that the driving forces for phase separation are stronger for −12F+12Y when compared to WT. However, what remains unresolved is whether the differences in driving forces are due to differences in effective protein-protein interactions, with the contributions of solvent components being equivalent, or if the replacement of Phe by Tyr alters the interplay of protein-protein, protein-solvent, and solvent-solvent interactions. For example, the lowering of *c*_sat_ for −12F+12Y, which would be implicitly attributed to enhanced protein-protein interactions, may be the result of solvent contributions that alter the nature and strengths of three-body interactions without influencing protein-protein associations [13]. Likewise, weakened protein-solvent interactions and enhanced solvent-solvent interactions, without any substantive changes to protein-protein interactions, could also account for changes in *c*_sat_. Being able to resolve the origins of changes to driving forces for phase separation requires knowledge of the extent to which different components contribute toward the driving forces for phase separation.

The problem of dissecting driving forces for phase separation becomes even more important in the context of living cells. Here, condensates comprise hundreds of different proteins and nucleic acids [1, 14, 15]. Further, the cellular milieu is a highly complex, non-ideal, osmotic solution, made up of an assortment of ions, osmolytes, and metabolites [16]. Additional complexities arise from the effects of macromolecular crowding and active processes within cells [17]. How does one deduce which components are important for phase separation? Riback et al. probed the effects of titrating the concentration of nucleophosmin (NPM1) on the biogenesis of nucleoli [18] in live cells. They estimated the apparent saturation concentration *c*_sat_ for NPM1, above which phase separation occurs. However, as the total NPM1 level increases beyond *c*_sat_, the concentration of NPM1 in the nucleoplasm, which is the dilute phase that coexists with nucleoli, does not stay fixed at *c*_sat_. Instead, it increases monotonically as the total concentration of NPM1 increases. The absence of saturating or plateauing behaviour suggests that components other than and in addition to NPM1 contribute to the assembly of the granular components of nucleoli. Riback et al. analysed this feature to show how a blend of homotypic (NPM1-NPM1) and heterotypic interactions (NPM1 interacting with other molecules) contribute to the assembly of facsimiles of granular components of nucleoli. The key quantity they studied, the partition coefficient of NPM1 between dilute and dense phases, encodes the inter-phase NPM1 chemical potential difference. This is however not the thermodynamic driver of phase separation. Instead, the onset of phase separation is governed by the total free energy difference difference between the transient, mixed state and the stable, phase-separated state. We refer to this difference as stabilisation free energy. In this work, we instead ask how can one infer the extent to which a chosen component contributes to the stabilisation free energy.

What is currently lacking is an unbiased formalism for assessing the dominant energetic contributions that only leverages the principles of phase equilibria without other assumptions. Here, we introduce a general framework that ties together the stabilisation free energy, the phase boundary and tie lines, and the experimentally accessible dilute phase concentration information (Fig 1). We consider a system comprising an arbitrary number *N* of different solute components, where two coexisting macro-phases viz., a macromolecule-rich and macromolecule-poor phase, result from phase separation. We do not assume any specific form of the free energy. Instead, we ask what information we can gain by analysing experimental data. Near the dilute phase boundary, we find that the stabilisation free energy associated with one solute relative to the whole system is closely linked to the shape of the phase boundary, slopes of tie lines, and dilute phase solute concentrations. Accordingly, we define the dominance of a solute as this stabilisation free energy fraction. We first demonstrate the flexibility and utility of the dominance parameter by analysing data from published simulations of 2-component systems. By comparing the dominance trend with those generated from a Flory-Huggins model we deduce the hierarchy of interaction strengths that are consistent with the data. In the second half of the work, we show that a simple dominance measurement approach exists for multi-component cases that can be adapted to any system of interest. In addition, we outline practical interpretations of dominance values in the context of interactions that modulate the driving forces for phase separation. The dominance framework is a new approach for quantitative investigations of phase separation systems.

**FIG. 1.**
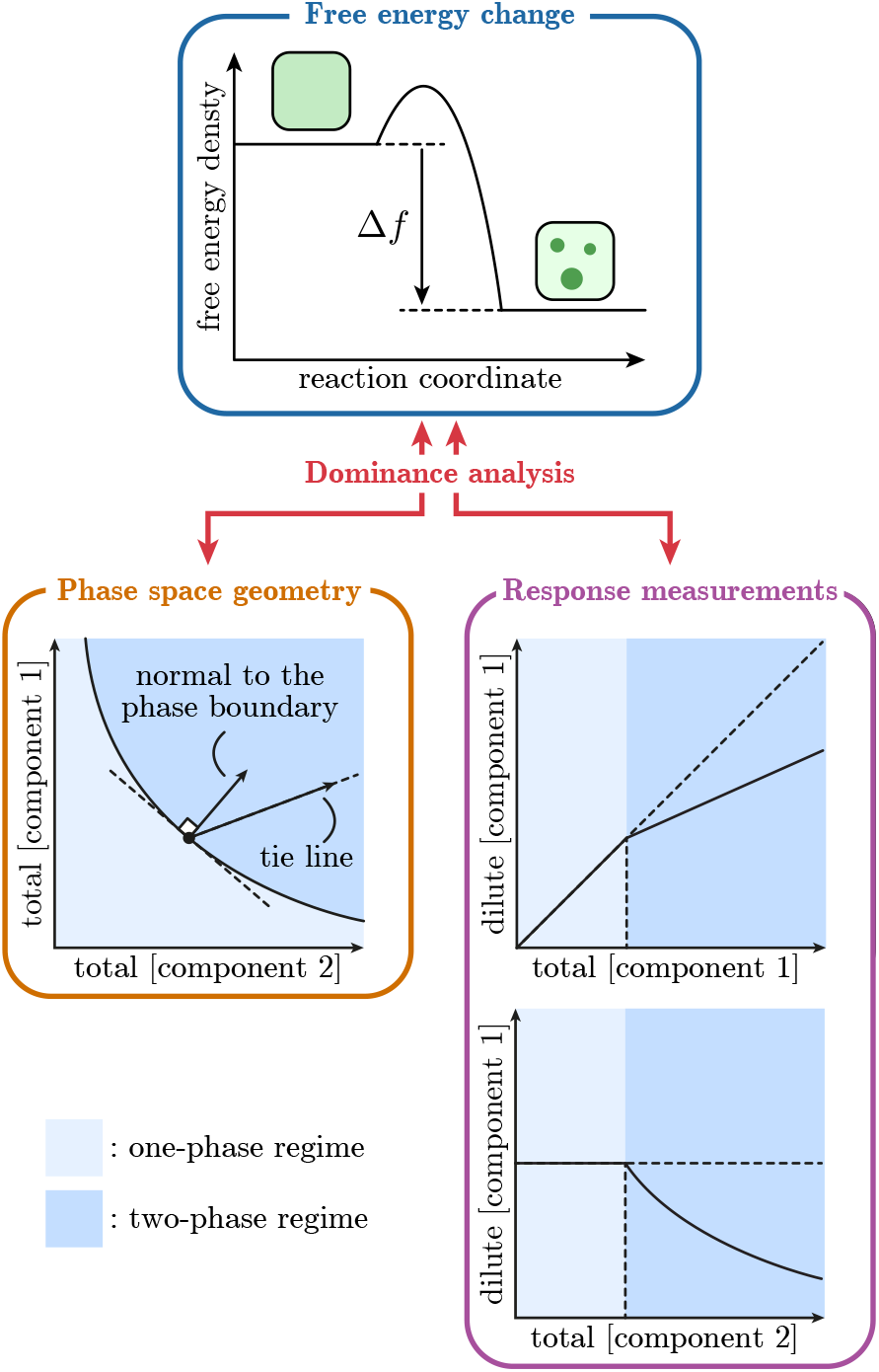
Overview of the dominance framework. The free energy change associated with phase separation Δ*f* is analysed in a model-free manner, and the relative contributions from different solutes towards Δ*f* is their respective dominance measure. Connections between dominance, the phase space, and dilute phase response functions are established in a unified framework.

## Defining dominance as an energetic measure

We denote the total concentrations, or equivalently volume fractions of solutes in a sample as *ϕ*^*α*^ where *α* = 1, 2,…, *N*. The dilute and dense phase boundaries can be represented by two (*N* − 1)-dimensional surfaces in the *N*-dimensional concentration space, and we use 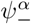 and 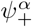 to denote solute concentrations on the dilute and dense surfaces respectively. Mass balance requires that for each sample *ϕ*^*α*^, the line connecting 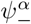 and 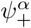 at equilibrium passes through *ϕ^α^*. We define the tie line vector 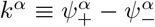 and use *v* to denote the volume of the dense phase relative to the whole system. The mass balance equation can then be written as 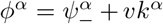. Furthermore, we use *n*_*α*_ to denote the vector normal to the dilute phase boundary, defined by 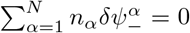 where 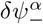 is a vector lying on the dilute arm of the phase boundary. This construction of the phase space is entirely geometrical and the only assumption is that two phases co-exist at equilibrium (Fig 2a). To fully define the space, we, in principle, need to assign a metric *g*^*αβ*^ that connects the units of different solute concentrations, and a natural choice is to use the volume fraction of solutes as a fundamental measure. This in turn allows a ‘distance’ to be defined, but neither the metric nor the distance is relevant for connecting the thermodynamics to experimental observables.

**FIG. 2.**
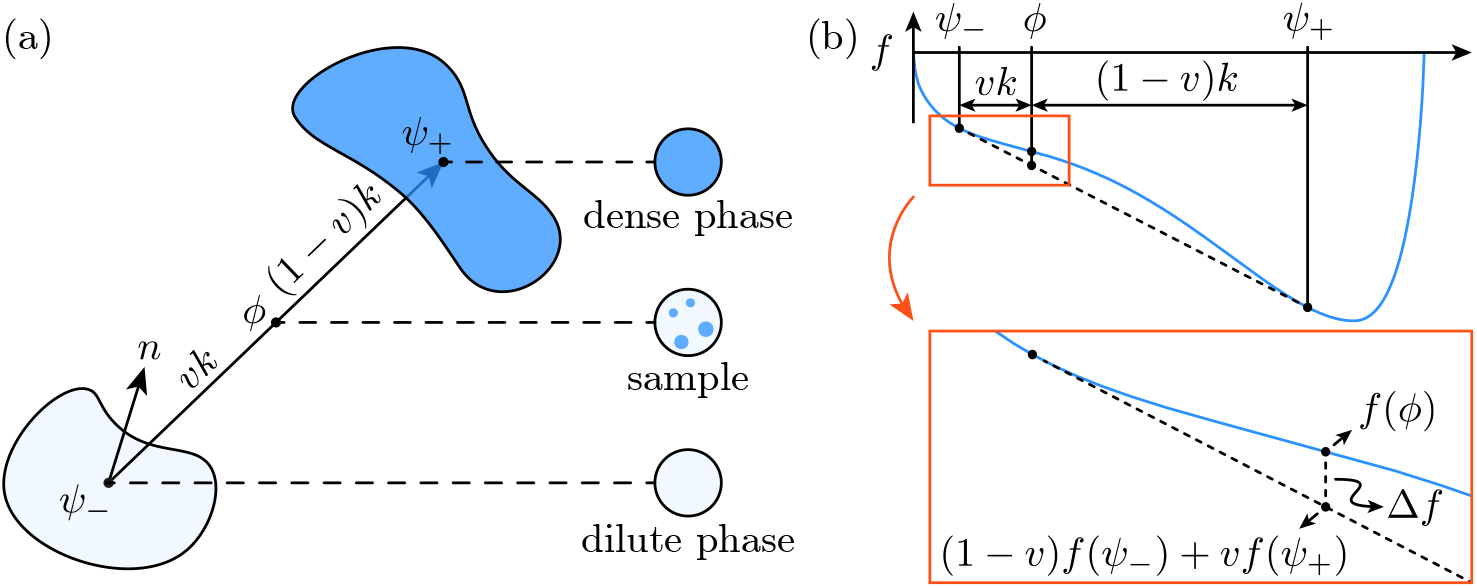
(a) The full *N*-dimensional phase space consists of dilute (light blue) and dense (dark blue) phase binodal boundaries, tie lines *k*^*α*^ connecting them, and dilute boundary surface normals *n*_*α*_. (b) The free energy before and after phase separation is *f* (*ϕ*) and (1 − *v*)*f* (*ψ*_−_) + *vf* (*ψ*_+_) respectively, and a 1-dimensional representation is used here but this is generalisable to any number of dimensions.

Phase separation occurs because it minimises the total free energy of the system. We denote this free energy density difference by Δ*f <* 0 (Figure 2b). We write the free energy density *f* (*ϕ*) of a system as a function of volume fractions of solutes *ϕ*^*α*^, and since the volume is constrained in defining *ϕ*^*α*^, *f* (*ϕ*) corresponds to the Helmholtz free energy. Under the assumption that the volume occupied by a molecule is fixed, it can be shown that *f* (*ϕ*) also plays the role of the Gibbs free energy density, so the results we obtained can be applied to the practical situation where the pressure, instead of volume, is held constant (SI section I). We compute Δ*f* in the limit of small *v*. This is equivalent to probing regions close to the dilute phase boundary. Δ*f* is given by (SI section II)

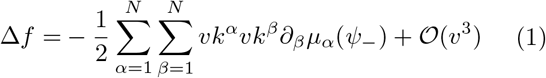

where the chemical potential is 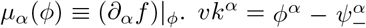 represents the displacement of the total composition from the dilute composition, so the sum over *β* can be viewed as computing the change in chemical potential of solute *α* upon phase separation. Accordingly, we define the individual term

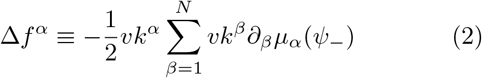

and it quantifies the free energy change associated with the partitioning of solute *α*. We are interested in 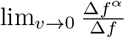, which quantifies the relative energetic contribution of solute *α* at the onset of phase separation. This leads to a natural definition of the dominance of *α* as

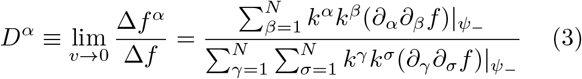

such that for each point 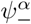 on the dilute phase boundary a *D*^*α*^ can be calculated. Notice that *D*^*α*^ is dimensionless and that the sum over all solutes is 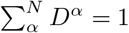. We are now in a position to deduce the relative ‘importance’ of a solute compared to others, thus the name ‘dominance’. Intuitively, if *D*^*α*^ for a solute *α* is found to be close to 1, this implies 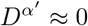 for *α*′ ≠ *α* and the phase behaviour is dominated by interactions involving the component *α* even through other components are present in the system. In general, for a multi-component system, we expect *D*^*α*^ to be in the range of (0,1) for more than one solute. *D*^*α*^ can thus serve as a way to identify and quantify the number of components that contribute to the dominant interactions that drive phase separation in multi-component system.

## Dominance computation from phase diagrams

Having formulated the dominance *D*^*α*^ we next explore how it can be computed from experimental observables. We motivate the following derivation by analysing simulation data for a system consisting of Low-Complexity Domains of protein Fused in Sarcoma (FUS-LCD) and hnRNPA1 (A1-LCD) [19]. Both FUS-LCD and A1-LCD undergo phase separation on their own, and by mixing them at different ratios while keeping their total concentration the same, a 2-dimensional phase diagram with tie lines can be produced [19]. Ratios between *n*_*α*_’s and *k*^*α*^’s can thus be computed using the gradient of the phase boundary and tie lines. The simulation results were reproduced *in vitro*, revealing strong heterotypic attraction between these proteins [19]. The simulation represents an exact 2-dimensional system, and to obtain estimates for the dominance values for the polymers we note that the tie line vector *k*^*α*^ already appears in Eq. (1), so we focus on the phase boundary normal *n*_*α*_. By perturbing the equilibrium compositions we arrive at 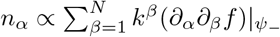 (SI section III) and direct substitution and comparison with Eq. (3) establishes the relationship between *D*^*α*^ and the geometrical quantities

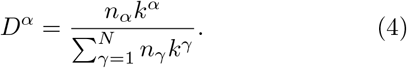

In the case of the 2-dimensional system [19] we have *α* = 1, 2, and we use index 1 to denote A1-LCD and 2 for FUS-LCD. The tie line gradient *k*^1^*/k*^2^ and the gradient of the phase boundary − *n*_2_*/n*_1_ can be combined to give the ratio *D*^1^*/D*^2^ = (*n*_1_*/n*_2_)(*k*^1^*/k*^2^). Using *D*^1^ + *D*^2^ = 1 we obtain individual estimates of *D*^1^ and *D*^2^ = 1 − *D*^1^. The simulations [19] were performed with a constant total concentration of A1-LCD and FUS-LCD, and 5 samples were simulated with varying percentages of each protein. Plotting the dominance of A1-LCD as a function of the percentage of FUS-LCD shows that when both proteins are present in the system, the dominance of A1-LCD is consistently larger than that of FUS-LCD (Figure 3, scatter points and dashed lines), and the dominance values are by definition 0 or 1 when either protein is absent from the system.

**FIG. 3.**
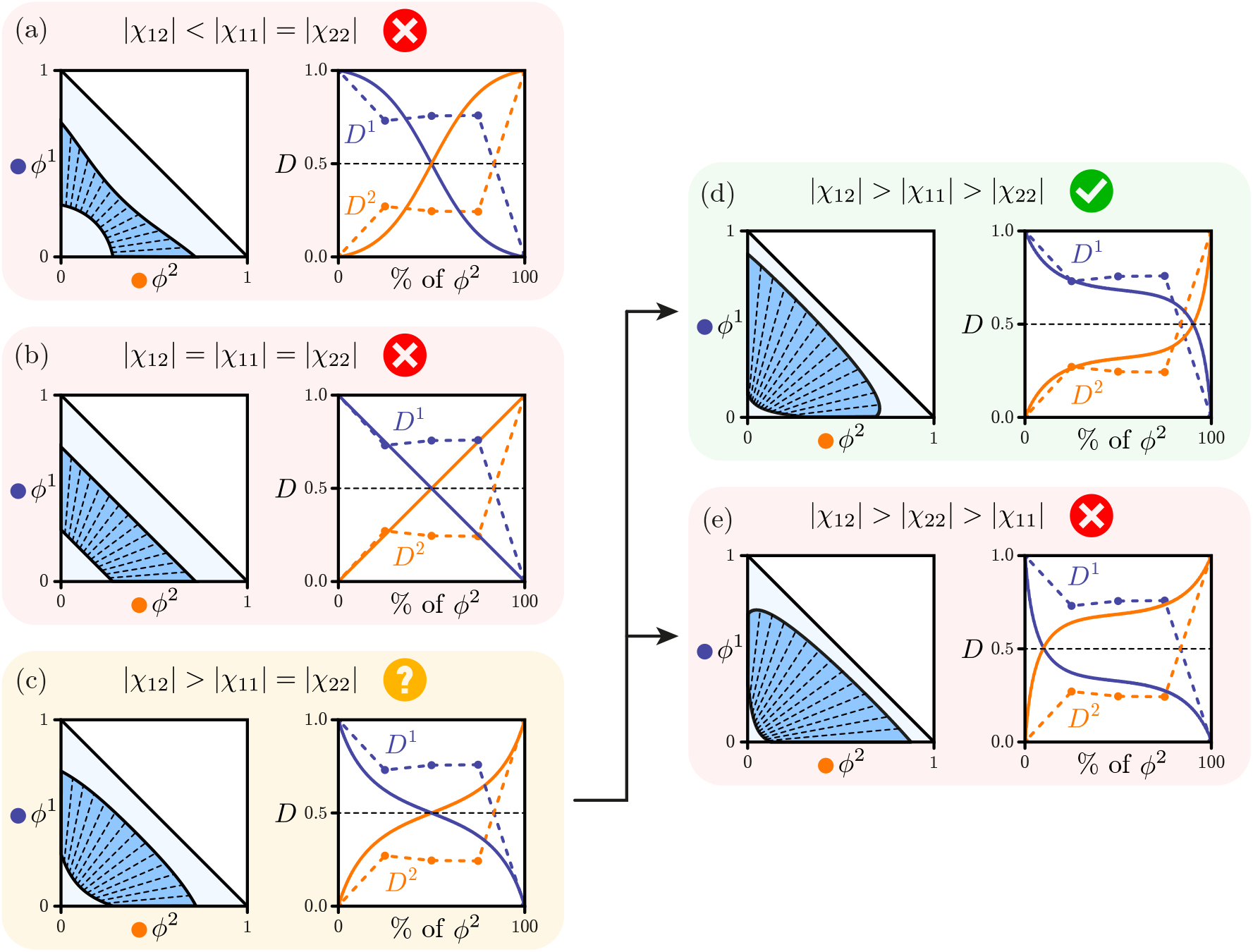
Computationally generated 2-dimensional phase diagrams from the Flory-Huggins model (left panels) and dominance values of the two solutes (right panels). Dominance values calculated from simulations in [19] are scatter points connected by dashed lines. In (a), (b) and (c) the homotypic interactions are the same, and the heterotypic interaction is weaker than, the same as, or stronger than the homotypic interactions respectively. Comparing the dominance trend to the A1/FUS system suggests the heterotypic interaction is likely stronger than homotypic attractions, and by further varying one of the homotypic interactions in (d) and (e) we deduce the homotypic interaction among *ϕ*^1^ is stronger than that among *ϕ*^2^.

To understand how relative interaction strengths in the simulation system give rise to the calculated dominance values, we used a 2-component Flory-Huggins model [5, 6] to generate the full phase diagram of a simplified system through the convex hull algorithm [20–22] (Appendix IV). In our computation we assume the two solutes are of unit length, fix *ϕ*^1^ + *ϕ*^2^ to be 0.5, and compute their dominance parameters as a function of the percentage of *ϕ*^2^. All interaction parameters are negative, so both solutes attract each other. We first fix the homotypic interactions *χ*_11_ and *χ*_22_ to be the same and vary the heterotypic interaction *χ*_12_ to be weaker than (Figure 3a), the same as (Figure 3b), or stronger than (Figure 3c) homotypic interactions. The dominance values for the former two cases vary rapidly as the percentage of *ϕ*^2^ increases. This trend is different from that in the data, so we deduce the heterotypic interaction is stronger than the homotypic interactions, which is in accord with published results [19]. To further estimate the relative strengths of homotypic interactions, we set *χ*_11_ to be stronger (Figure 3d) or weaker (Figure 3e) than *χ*_22_. The dominance trend produced in the former case is qualitatively similar to the simulation data. We conclude that not only is the heterotypic attraction the strongest, the homotypic attraction among A1-LCD is stronger than that among FUS-LCD. It is worth noting that the dilute phase boundary from simulations is orders of magnitude smaller than in the Flory-Huggins computation, while the dimensionless nature of *D*^*α*^ means this difference can be ignored and only important features of the phase space, including the phase boundary gradient and the tie lines, are relevant.

Next, we study a very different scenario where phase separation is driven purely by the heterotypic interaction. We set χ_11_ = χ_22_ = 0 and as established before [23], phase-separation can happen with either strong heterotypic attraction (Figure 4a) or heterotypic repulsion (Figure 4b) in the absence of any homotypic interaction. In presenting dominance values we also use a different parameterisation: for each point on the dilute phase boundary we compute the angle *θ* made between the vector connecting the origin to that point and the *ϕ*^1^ axis, and plot *D*^*α*^ as a function of *θ*. This is qualitatively the same as plotting *D*^*α*^ as a function of the percentage of *ϕ*^2^ except in the case of repulsive interactions, where the negative tie line gradient means *D*^*α*^ stay constant if the total solute concentration is fixed. Repeating the *D*^*α*^ calculation now gives drastically different results. With strong heterotypic attraction (Figure 4a), the computed *D*^1^ starts off at a small value and increases as more *ϕ*^2^ is added. This effect is of entropic origin: at low *ϕ*^2^, the translational entropy of *ϕ*^2^ opposes phase-separation, and as *ϕ*^2^ increases this opposition diminishes due to the logarithmic scaling of the entropic free energy. As a result, the interaction free energy previously counterbalanced by the entropic free energy is released. Consequently, the free energy release by *ϕ*^2^ is more significant compared to that by *ϕ*^1^ and thus *D*^2^ attains a much larger value than *D*^1^. Conceptually, the dominance also reflects which solute is ‘limiting’ in the phase separation process. With heterotypic repulsion (Figure 4b), defining dense and dilute phases is non-trivial because the solutes partition into separate phases. Here we focus on the region of phase space where *θ > π/*4 so that the dense phase is higher in *ϕ*^1^. In this region, *D*^1^ *>* 1 throughout and this implies *D*^2^ = 1 − *D*^1^ *<* 0. This can appear counterintuitive, as it appears that *ϕ*^2^ is in an energetically less favourable state after phase-separation. This is explained by the fact that by phase-separating, the change in chemical potential of *ϕ*^1^ due to partitioning of *ϕ*^2^ releases free energy to balance out the unfavourable contribution from *ϕ*^2^ itself, and thus *ϕ*^2^ becomes a depletent. This behaviour is expected of real crowders [24, 25] and it has been reported recently by Chauhan et al. [26] for the phase behaviour of transcription factors in the presence of PEG. It is also worth noting that in both cases, as the system moves towards a critical point, *D*^1^ appears to approach a singularity and this arises because at the critical point, Δ*f* = 0 and the dominance becomes ill-defined.

**FIG. 4.**
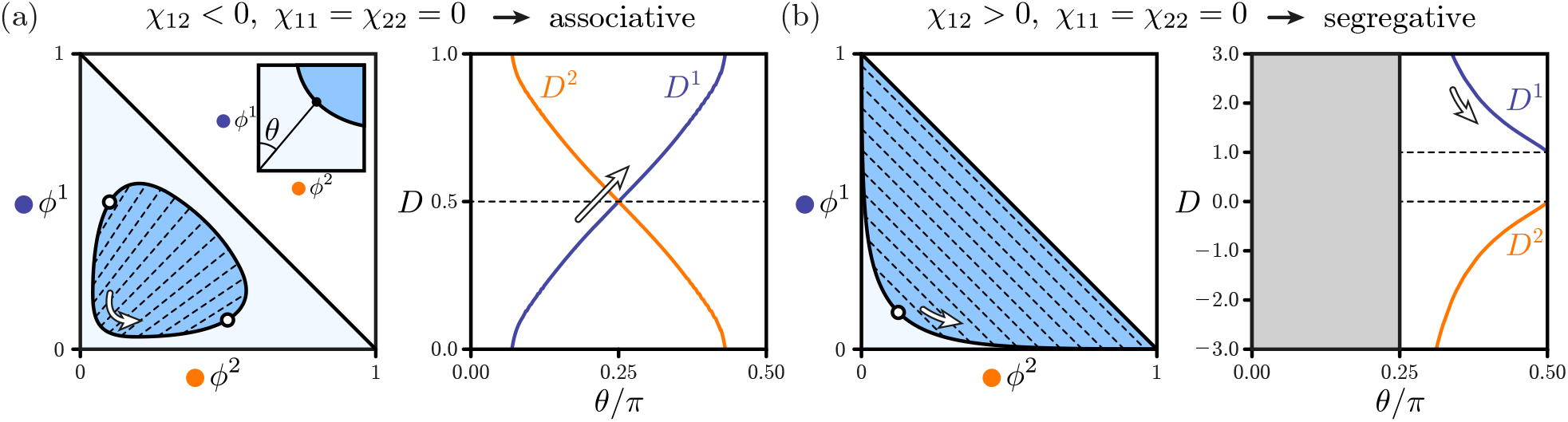
When phase-separation is driven purely by heterotypic interactions, both attraction (e) and repulsion (b) can lead to de-mixing. Left panels show the 2-dimensional phase diagrams with white points as critical points, and computed *D*^*α*^ values are plotted in the right panels. Inset: *θ* is defined as the angle formed between the dilute phase point where the dominance is evaluated, and the *ϕ*^1^ axis. As the point of evaluation approaches the critical points (black arrows) the *D*^*α*^ values encounter a singularity as Δ*f* goes to 0. An interesting observation in (b) is that here, *D*^1^ takes on values not in the range of (0, 1), reflecting the abnormal behaviour of a repulsion-driven phase-separating system.

Motivated by the computational models, we outline insights that can be gained if the dilute phase concentrations regarding multiple solutes can be measured. If *D*^*α*^ and ratios of *n*_*α*_ and *k*^*α*^ can be measured at each experimental point, entries in the Hessian ∂_*α*_∂_*β*_*f* (*ψ*_−_) can be obtained as well, and these correspond to interaction energies of solutes in the dilute phase. This is based on the observation that 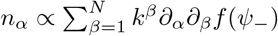. To illustrate this point, suppose ∂_*α*_∂_*β*_*f* (*ψ*_−_) consists of two terms: a constant, symmetric interaction energy matrix χ_*αβ*_ and a diagonal entropic term 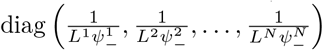 where *L*^*α*^ are sizes of the solute molecules. In doing so the free energy from translational entropy of solute *α* is assumed to be 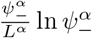. Treating the interaction matrix and sizes *L*^*α*^ as unknowns, we have 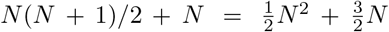 fitting parameters.

On the other hand, at each point 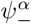, we have *N*− 1 linearly independent equations using ratios of *n*_*α*_’s. For an *N*-component system by sampling 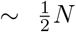 points, the interaction constants χ _*αβ*_ and molecular sizes *L*^*α*^ can be deduced. Practical implementation of this procedure can however be difficult, given the large number of relevant solutes. Nonetheless, the prospect of obtaining *χ*_*αβ*_ in the dilute solution not only helps with identifying the role of different solutes to phase separation, but also quantifies protein interactions in general.

## Dilute phase response function

The 2-dimensional systems investigated so far serve as illustrative examples of the dominance formulation, and we now turn to the realistic scenario where the number of solute species can be more than two. In addition, concentrations of many of these species can be hard to quantify, so full phase diagrams and tie lines are rarely accessible. We observe, however, that in a typical experiment there will be at least one macromolecular solute, say *α*, whose dilute phase concentration is quantifiable via fluorescent labelling and microscopy methods [11, 23]. Furthermore, one can also change the total starting concentrations of any solute *β*, so an accessible quantity of interest is the response 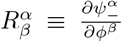. Via mass conservation, one can show that changes in 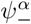 in response to changes in *ϕ*^*β*^, evaluated at the phase boundary, can be expressed as (SI section V)

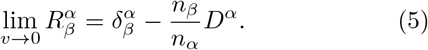

The appearance of *D*^*α*^ indicates that only the dominance of *α* is accessible when the dilute phase concentration of *α* is measurable, regardless of which solute *β* is varied to measure the response. Interestingly, by combining two response functions we can further deduce information regarding the tie line components *k*^*α*^ and *k*^*β*^.

Previously [23] we established a tie line measurement approach between a solute with index 1 (without loss of generality), and another solute with index 2 through measurement of 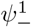 alone. Using the response function formulation we can ask the following question: how should *ϕ*^1^ and *ϕ*^2^ be changed so that 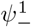 stays constant? Mathematically, this requires 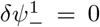. Recall that 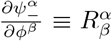, and 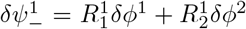. The space defined by 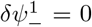 on the 1 − 2 plane is a line, and its gradient is 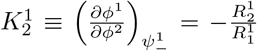. Close to the dilute phase boundary we can use Eqs. (4) and (5) to re-write 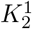 as

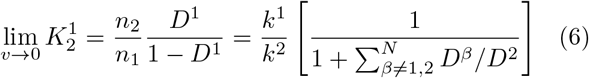

such that to leading order, 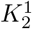 provides an approximation of the ratio of tie line components 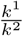 close to the phase boundary. Terms in the square bracket of Eq. (6) represent the relative deviation of 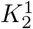 from 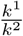, and in a true 2-component system we have 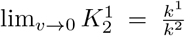. On the 1 − 2 plane, this line defined by 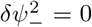 can be thought of as an *N*-dimensional tie line ‘reduced’ to the 2-dimensional plane, and we refer to it as the reduced tie line in the following discussions.

Measuring the gradient of the reduced tie line allows an estimate of the relative solute partitioning to be made. Notably, if the dominance of all solutes falls within the range (0, 1), the sign of 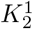 is entirely determined by the sign of 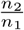, and thus the sign of 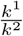 can be deduced by observing the effect of the solute 2 on condensates. To put this in practical terms, if solute 2 dissolves Condensates 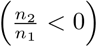 it partitions out of condensates 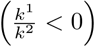 and vice versa. This makes intuitive sense since it suggests that condensate-favouring modulators partition into condensates while condensate-disfavouring modulators are excluded from condensates. It is important however to point out here that some *D*^1^ can fall outside the 0 and 1 range, and this can happen when a solute is able to change the chemical potentials of other solutes to make phase separation especially favourable, even though its own energetic contribution is opposing phase separation. This is the case of true crowders, as explained in the previous section.

## Response measurement using line-scans

We now investigate practical applications of dominance measurements and interpretations in a range of systems. To obtain the dominance of a solute from the dilute phase responses it is necessary to measure its dilute phase concentration and in the following we use the index 1 to denote the solute that is measured. Experimentally, a series of samples can be prepared with varying *ϕ*^1^ (homotypic line-scan) or some other *ϕ*^2^ (heterotypic line-scan) while keeping all other *ϕ*’s constant and measuring 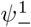. This corresponds to traversing the phase space along a line parallel to *ϕ*^1^ or *ϕ*^2^ axis and measuring the response functions 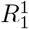 or 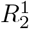. By performing the measurement close to the dilute phase boundary it is convenient to rewrite response expressions:

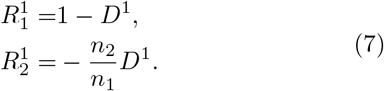

We use published experimental data on phase separation of the optoG3BP1 protein in cells [18] where the dilute phase [optoG3BP1] was measured as a function of total [optoG3BP1], and use the index 1 to denote the optoG3BP1 protein. We have 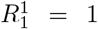 outside the phase-separated region, and the response becomes modulated beyond some threshold value 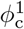. This is in essence the apparent saturation concentration *c*_sat_ in other literature. In an ideal 1-dimensional system, for 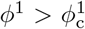 we expect 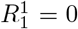. Instead, we find 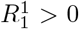 for optoG3BP1 because of the multi-component nature of the system [18] (Figure 5a). Using a 2-segment linear fit for optoG3BP1 data we obtain *D*^1^ = 0.55 ± 0.14. In contrast, in another system involving the protein opto-FUS [27], the dominance of optoFUS is *D*^1^ = 0.98± 0.07 (Figure 5b). This is consistent with the observation that FUS phase separates on its own [29, 30] and can be less reliant on other components compared to G3BP1, where other macromolecules such as RNA are typically added to G3BP1 solution to trigger phase separation [31, 32]. A simple dominance measurement thus allows a quick assessment of the multi-component character, similar to what Guillen-Boixet et al., demonstrated [31]. Taking a step further, we explore how the system response to addition of modulators can be studied using the energy dominance analysis.

**FIG. 5.**
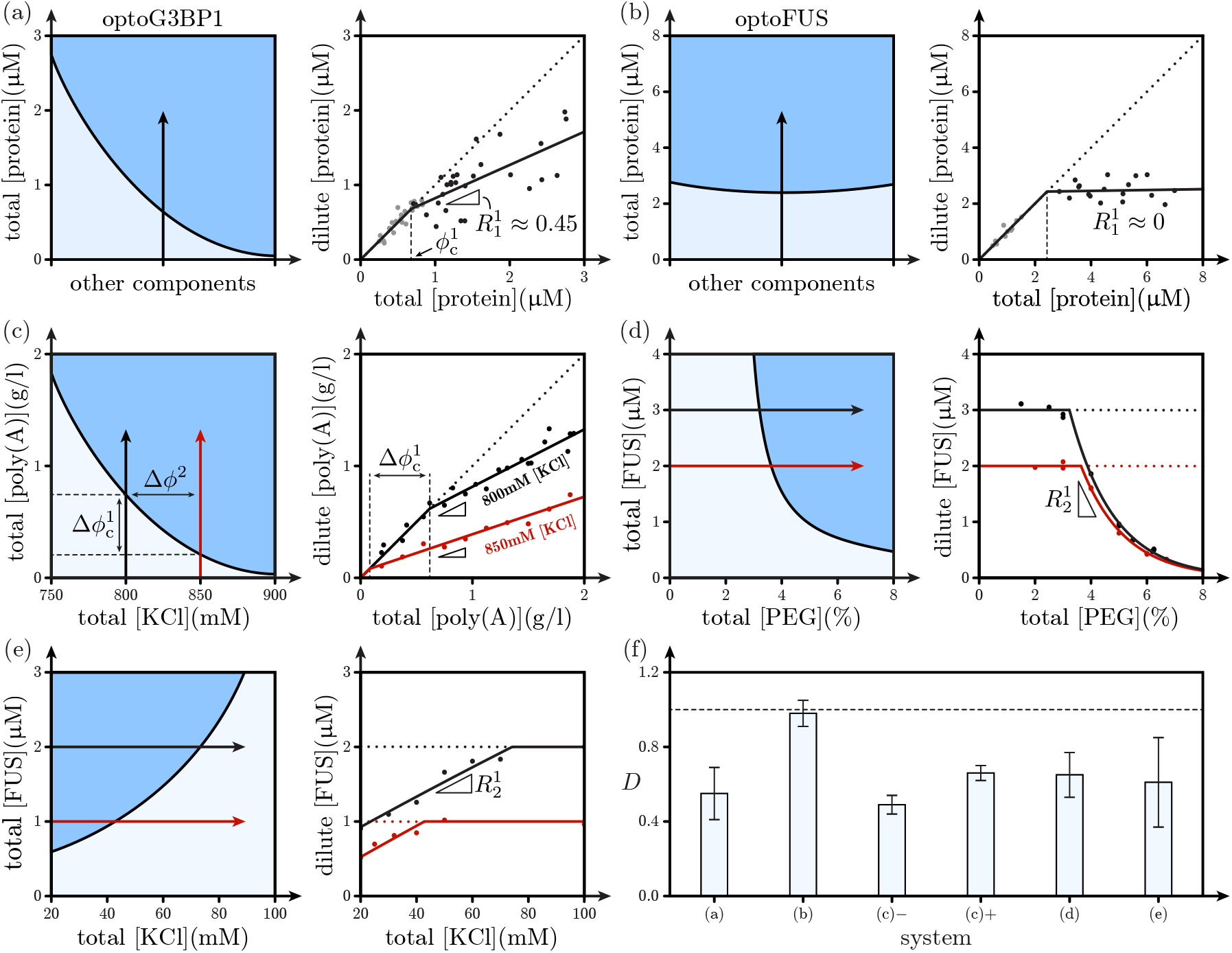
In (a) to (e), right panels illustrate cross-sections of phase spaces, left panels are data. Arrows in illustrations represent how line-scans are traversing the phase spaces. In data panels, scatter points are data and solid lines are phenomenological fits. Dotted lines indicate trivial results in the absence of phase-separation. (a) The dilute phase response of optoG3BP1 [18] exhibits multi-component character, so phase-separation is likely to depend on other components. (b) In the optoFUS system[27]the protein has much stronger single-component character. (c) Poly(A)-RNA phase-separates in the presence of PEG and KCl. At higher [KCl] the increased charge screening allows inter-RNA interaction to be more favourable, leading to an increase in *D*^1^ when [KCl] is increased. (d) Heterotypic line-scan of FUS against PEG [23]. At low [PEG] no phase separation is observed, so the dilute phase [FUS] is simply a constant. When the line-scan trajectory enters the phase separated region the dilute phase [FUS] starts decreasing, and this initial gradient is taken as 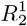. (e) Heterotypic line-scan of FUS against KCl [28]. Here the plateau appears at high [KCl] because KCl dissolves condensates. (f) Overview of dominance values collected. − and + labels for (c) correspond to low and high [KCl] respectively.

Poly(A)-RNA phase separates in the presence of PEG and at high [KCl] [33, 34], and here we investigate how the dominance of poly(A) changes upon addition of salt. We perform homotypic line-scans with 4% PEG (molecular weight 10 kilodaltons), and 800 mM or 850 mM of KCl (SI section VI). Dilute phase poly(A) concentrations were measured, so we use index 1 to denote poly(A) and index 2 for KCl, and denote the difference in total KCl concentrations as Δ*ϕ*^2^ = 50 mM. Line-scan data show a smaller 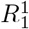 after phase-separation at higher KCl so *D*^1^ increases from 0.49 ± 0.05 to 0.66 ± 0.04 (Figure 5c). The increase in poly(A) dominance at higher KCl can be explained by the increased screening of negative charges on poly(A), facilitating poly(A)/poly(A) interaction and rendering other interactions, for instance poly(A)/PEG, less significant. An interesting observation is that the onset of phase-separation shifts to a lower poly(A) concentration when salt concentration is increased, with 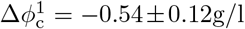. This can be interpreted as an increase in overall phase-separation propensity, consistent with the picture that poly(A)/poly(A) repulsion weakens with more salt. By combining Δ*ϕ*^2^ with 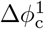 we can deduce the ratio of surface normal 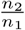 in the poly(A)-KCl plane. By definition, 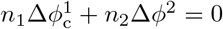 so

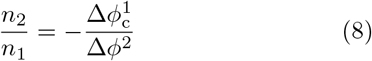

and this gives 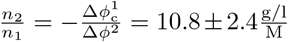. By combining this with the average *D*^1^, we can further calculate the gradient of the reduced tie line as 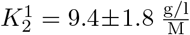. The simple inference is that KCl partitions preferentially into poly(A) condensates. However, in reality we expect the inter-phase partitioning of K^+^ and Cl− to be different and dependent on the total concentration of KCl, and this information can only be gained by measurements of individual ionic species. Mathematically, the line-scan does not traverse parallel to one axis but at an angle between two axes corresponding to K^+^ and Cl−, so mixing them is equivalent to performing a rotation of the two axes. This does not affect 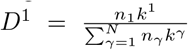 since the denominator is coordinate-independent and the numerator is not affected. Furthermore, since *k*^*α*^ and *ϕ*^*α*^ transform in the same way the measured 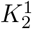 represents the summed partitioning of K^+^ and Cl−, but it is blind to individual partitioning.

The homotypic line-scan is a natural path towards quantifying *D*^1^, while in experiments the heterotypic line-scan can also be performed [23, 28]. To illustrate the approach, we take two sets of data: FUS/PEG from [23] and FUS/KCl from [28]. In both cases dilute phase FUS protein (index 1) concentration is measured, while the total concentrations of PEG or KCl (index 2) are varied. In the FUS/PEG line-scan data from [23], two line-scans were performed at 2 and 3 μM [FUS] while adding PEG up to 8% weight fraction. The dilute phase [FUS] measurements showed a plateau at low [PEG] and an exponential decrease as some thresholding [PEG] is reached (Figure 5d). The two points at the onset of the decrease are where the lines cross the phase boundary, and from these we estimate 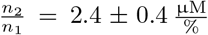. The response function 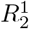 can be evaluated at the drop-off points as well, averaging over the two scans gives 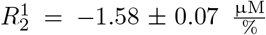. Combining 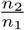 and 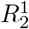 gives *D*^1^ = 0.65 ± 0.12. This dominance value indicates there are other solutes actively participating in phase-separation and a likely candidate is PEG. This is also reflected in the fact that the FUS/PEG phase boundary measured using PhaseScan has strong dependence on [PEG] [35]. Furthermore, substituting 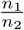 and the average *D*^1^ into Eq. (6) we get 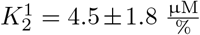, providing an estimate of the relative partitioning between FUS and PEG. In the FUS/KCl system [28], KCl has the opposite effect compared to PEG as condensates are dissolved when more KCl is added to the system [28]. Line-scan data thus shows a steady increase of dilute phase [FUS] with increasing [KCl] (Figure 5e). We fit the dilute phase [FUS] after phase-separation using a linear function. Repeating the same calculations we obtain 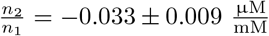 and 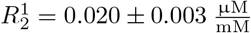 to give *D*^1^ = 0.61 *±* 0.24. This again suggests a FUS-dominated, multi-component system. The 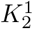 value obtained is 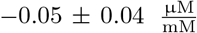, implying overall salt is partitioning outside condensates. All dominance values obtained are summarised in a bar plot (Figure 5f). Heterotypic line-scans are thus an alternative to homotypic line-scans, while the two methods revolve around the same theme of dominance analysis.

## Conclusion

We formulated a dimensionless parameter *D*^*α*^ that can be used to discern the extent to which a solute *α* contributes to the driving forces for phase separation in a multicomponent system. Our analysis focused on phase separation that gives rise to two coexisting phases. We demonstrated the accuracy and flexibility of the dominance framework by deploying it to analyse simulation results for mixtures of low complexity domains as well as experimentally measured dilute phase response functions.

Condensates such as stress granules are multicomponent mesoscale entities. Yang et al., used systematic knockdown experiments to converge on G3BP1/2 as the central node in the network of proteins that make up stress granules [32]. Sanders et al., showed that different nodes in the network contribute differently to the G3BP1/2-mediated assembly of stress granules [36], which is potentiated by stress-induced polysomal runoff that generated protein-free, unfolded RNA molecules [31]. As proteome-scale investigations become more widely used [37–39], it becomes increasingly feasible to identify nodes in protein-protein and protein-nucleic acid interaction networks that are key contributors to condensate assembly. Assessing the contributions of different condensate components will likely involve genetic manipulations of expression levels that enable homotypic and/or heterotypic scanning [18, 36, 37, 40–42]. However, the number of components within the condensate will always vastly exceed the number of independent axes along which solute concentrations can be titrated within cells [43]. Deployment of dominance analysis introduced in this work, and ongoing generalisations of this analysis will help quantify the contributions of titratable versus non-titratable and hence hidden components to the driving forces for condensation.

Here, we have shown that quantitative estimates of the solute-specific dominance parameters allow for insights to be gleaned regarding components that drive phase separation versus components that function as modulators. Recent studies adapted the polyphasic linkage formalism of Wyman and Gill [44] to understand how ligands that bind site-specifically can exert an influence, via linkage between binding and phase equilibria, on the driving forces for phase separation [45–47]. The formalism of binding polynomials, which underlies linkage relationships, is also free of any assumptions regarding the models used to describe either phase separation or binding. Combining the formalism of polyphasic linkage, which rests on quantifying how ligands modulate *c*_sat_, and dominance analysis, which dissects the energetic contributions of different solutes, might be a useful route for uncovering differences in binding modes across the phase boundary.

In multicomponent systems, the interplay between homotypic and heterotypic interactions determines the shapes of phase boundaries and the slopes of tie lines. Lin et al. showed that the shapes of phase boundaries, mapped in terms of concentrations of interaction motifs rather than molecules, can be used to dissect the relative contributions of homotypic versus heterotypic interactions [48]. A comparative assessment of the shape analyses introduced by Lin et al. and the dominance analysis introduced here would be valuable, especially for analyzing low-dimensional representations of phase diagrams for multicomponent systems, which are inherently multidimensional in nature. Other contributions including the concept of solubility products, which was recently introduced by Chattaraj et al., [49] the work of Deviri and Safran [50], both of which focused on buffering in the presence of heterotypic interactions, can be augmented by the dominance analysis introduced here.

Formally, the dimensionless nature of the dominance allows direct comparison between solutes that are distinct from one another. This makes it possible to compare the dominance of a macromolecular solute with that of a small molecule species, if the dilute phase concentration of the latter can be measured. Dominance analysis takes on special relevance given the likely contributions of metabolites, osmolytes, other naturally occurring small molecules, and drug-like molecules that are either functional regulators of condensation or are introduced to regulate condensate stability in vivo [51].

## Acknowledgement

This work was funded by grants from Global Research Technologies, Novo Nordisk A/S (H.A., T.P.J.K.), the European Research Council under the European Union’s Horizon 2020 research and innovation programme through the Marie Sklodowska-Curie grant MicroREvolution (agreement ID 101023060; T.S.), the Newman Foundation (T.S., T.P.J.K.), the European Research Council under the European Union’s Horizon 2020 research and innovation program through the ERC grant DiProPhys (agreement ID 101001615; T.P.J.K.), the US National Institutes of Health (R01NS121114 to R.V.P), and the US National Science Foundation (MCB-2227268 to R.V.P).

## Conflict of interests

The authors declare no conflict of interests.

## Notes

### Competing Interest Statement

The authors have declared no competing interest.

### Summary of Updates

Introduction updated; added summary figure 1; updated figure 5 layout

