## Supplementary Information for "Dominance Analysis: A formalism to uncover dominant energetic contributions to biomolecular condensate formation in multicomponent systems"

### SI for: Dominance formulation in multi-component binary phase equilibria

**Notations.** With the exception of SI section I, in this supplementary information document we follow conventions from differential geometry, where summation over repeated greek indices from 1 to $N$ is assumed unless otherwise stated. In cases where a summation is not implied, or if an index correspond to a scalar quantity instead of a vector component we simply put a round bracket around the index, e.g. $(\alpha)$. For instance the dominance $D^{(\alpha)}$ is coordinate-dependent [it depends on the alignment of the $(\alpha)$-axis relative to the phase boundary] and it is strictly speaking not a vector quantity. It is possible to define a metric tensor in this framework but we note that throughout the derivation, the metric plays no role as all units cancel out in the computation of $D^{(\alpha)}$.

#### I. EQUIVALENCY BETWEEN THE GIBBS AND HELMHOLTZ FREE ENERGIES

**Notations.** Notations in this section is separate from the rest of the SI due to many overlapping symbols, also because the derivations here are not relevant to the main theoretical results but serve to simply bridge the two free energies together.

In this section we explore how the Gibbs free energy is linked to the Helmholtz free energy under the assumption of zero compressibility. The Gibbs free energy $G$ of a system is a function of numbers of molecules of each species, including the solvent. We use the index 0 to denote the solvent molecule and $1, 2, \ldots, N$ for the solutes. Summations are also explicitly written out. Denote the number of molecules of each species as $n^\alpha$ the Gibbs free energy is a function of all $n^\alpha$:

$$G = G(n^0, n^1, n^2, \ldots, n^N) \tag{S.1}$$

and in writing so, no assumption about the volume is made, and $G$ may change as the pressure of the system is changed. We omit the pressure in the variables and assume it is held constant.

The equilibrium conditions when two phases form, each with number of molecules $n_-^\alpha$ and $n_+^\alpha$ respectively, are now simply

$$\left.\frac{\partial G}{\partial n^\alpha}\right|_{n_-} = \left.\frac{\partial G}{\partial n^\alpha}\right|_{n_+}, \quad \alpha = 0, 1, \ldots, N. \tag{S.2}$$

In the rest of this section we wish to show that by changing the variable from numbers $n^\alpha$ to volume fractions $\phi^\alpha$ we can construct a Helmholtz-like free energy density $f$ that re-casts Eqs. (S.2) into

$$\begin{aligned} \left.\frac{\partial f}{\partial \phi^\alpha}\right|_{\phi_-} &= \left.\frac{\partial f}{\partial \phi^\alpha}\right|_{\phi_+}, \quad \alpha = 0, 1, \ldots, N \\ \sum_{\alpha=1}^{N} \left.\frac{\partial f}{\partial \phi^\alpha}\right|_{\phi_-} \phi_-^\alpha - f(\phi_-) &= \sum_{\alpha=1}^{N} \left.\frac{\partial f}{\partial \phi^\alpha}\right|_{\phi_+} \phi_+^\alpha - f(\phi_+) \end{aligned} \tag{S.3}$$

which are those followed by the canonical Helmholtz free energy.

We denote the volume of one molecule of species $\alpha$ as $v^\alpha$, and assume these are constants. The total volume $V$ is

$$V = v^0 n^0 + \sum_{\alpha=1}^{N} v^\alpha n^\alpha \tag{S.4}$$

and we can re-write $n^0$ as

$$n^0 = \frac{V - \sum_{\alpha=1}^{N} v^\alpha n^\alpha}{v^0} \tag{S.5}$$

so that we can use $V$ as a variable that replaces $n^0$. We define the new function $G' = G'(V, n^1, n^2, \ldots, n^N)$ as

$$G'(V, n^1, n^2, \ldots, n^N) \equiv G\left(\frac{V - \sum_{\alpha=1}^{N} v^\alpha n^\alpha}{v^0}, n^1, n^2, \ldots, n^N\right). \tag{S.6}$$

Next, we can also define the volume fraction of solute $\alpha$ as $\phi^\alpha \equiv v^\alpha n^\alpha / V$ and use this to replace $n^\alpha$. Denote the new function as $G'' = G''(V, \phi^1, \phi^2, \ldots, \phi^N)$ we write

$$G''(V, \phi^1, \phi^2, \ldots, \phi^N) \equiv G'\left(V, \frac{V\phi^1}{v^1}, \frac{V\phi^2}{v^2}, \ldots, \frac{V\phi^N}{v^N}\right). \tag{S.7}$$

Eqs. (S.6) and (S.7) can be combined to give

$$G''(V, \phi^1, \phi^2, \ldots, \phi^N) = G\left(\frac{V - V\sum_{\alpha=1}^N \phi^\alpha}{v^0}, \frac{V\phi^1}{v^1}, \frac{V\phi^2}{v^2}, \ldots, \frac{V\phi^N}{v^N}\right). \tag{S.8}$$

Finally we define the free energy density $f = f(\phi^1, \phi^2, \ldots, \phi^N)$ by normalising $G''$ with respect to $V$:

$$f(\phi^1, \phi^2, \ldots, \phi^N) = \frac{G''(V, \phi^1, \phi^2, \ldots, \phi^N)}{V} \tag{S.9}$$

and the dependence of $V$ has been dropped in $f$ because it is easily verified that $\frac{\partial f}{\partial V} = 0$.

With the $f$ defined, we now show that Eqs. (S.3) will reproduce Eqs. (S.2). Calculating the chemical potential in the Helmholtz sense:

$$\frac{\partial f}{\partial \phi^\alpha} = \frac{1}{V}\frac{\partial G''}{\partial \phi^\alpha} = \frac{1}{V}\left[\frac{\partial G}{\partial n^0}\left(-\frac{V}{v^0}\right) + \frac{\partial G}{\partial n^\alpha}\left(\frac{V}{v^\alpha}\right)\right] = \frac{\mu^\alpha}{v^\alpha} - \frac{\mu^0}{v^0}, \quad \alpha = 1, 2, \ldots, N. \tag{S.10}$$

where $\mu^\alpha \equiv \frac{\partial G}{\partial n^\alpha}$ for $\alpha = 0, 1, 2, \ldots, N$. On the other hand, the 'osmotic pressure' is

$$\sum_{\alpha=1}^N \frac{\partial f}{\partial \phi^\alpha}\phi^\alpha - f = \sum_{\alpha=1}^N \left(\frac{\mu^\alpha}{v^\alpha} - \frac{\mu^0}{v^0}\right)\left(\frac{v^\alpha n^\alpha}{V}\right) - \frac{G}{V} = -\frac{\mu^0 n^0}{V} - \frac{\mu^0}{v^0}\frac{V - n^0 v^0}{V} = -\frac{\mu^0}{v^0} \tag{S.11}$$

where we have used $G = \sum_{\alpha=0}^N \mu^\alpha n^\alpha$ in the second equality. Substituting Eqs. (S.10) and (S.11) into Eqs. (S.3) then produces Eqs. (S.2).

#### II. STABILISATION ENERGY OF PHASE SEPARATION

The general principle of phase equilibria requires equilibration of chemical potentials and osmotic pressure across phases [1]. Formulating this process as a free energy minimisation problem constrained by global conservation of order parameters $\phi^\alpha$ [2, 3] leads to definitions of chemical potentials $\mu_\alpha(\phi) \equiv \frac{\partial f(\phi)}{\partial \phi^\alpha} \equiv \partial_\alpha f(\phi)$ and the osmotic pressure $\Pi(\phi) \equiv \phi^\alpha \mu_\alpha(\phi) - f(\phi)$, so that the resulting binary phase equilibria satisfies

$$(\partial_\alpha f)|_{\psi_-} = (\partial_\alpha f)|_{\psi_+}, \tag{S.12a}$$

$$\psi_-^\alpha (\partial_\alpha f)|_{\psi_-} - f(\psi_-) = \psi_+^\alpha (\partial_\alpha f)|_{\psi_+} - f(\psi_+), \tag{S.12b}$$

$$\tag{S.12c}$$

corresponding to conditions on chemical potentials and the osmotic pressure. The assumption of binary phase-separation is made implicit by assuming only two phases are present. The average free energy density change after phase separation is

$$\Delta f = (1 - v)f(\psi_-) + vf(\psi_+) - f(\phi). \tag{S.13}$$

To make progress, we substitute Eq. (S.12a) into Eq. (S.12b) to obtain $f(\psi_+) = f(\psi_-) + k^\alpha(\partial_\alpha f)_{\psi_-}$ where we have used $k^\alpha = \psi_+^\alpha - \psi_-^\alpha$. The mass balance $\phi^\alpha = \psi_-^\alpha + vk^\alpha$ can be used to expand $f(\phi)$ as a power series in $vk^\alpha$: $f(\phi) = \sum_{i=0}^\infty \frac{1}{i!}\left[(vk^\alpha \partial_\alpha)^i f\right]|_{\psi_-}$. By limiting the dense phase volume fraction $v$ to be small, equivalent to probing regions close to the dilute phase boundary, the sum is dominated by the first few terms. Substituting $f(\psi_+)$ and $f(\phi)$ into $\Delta f$ gives

$$\begin{aligned}
\Delta f =& (1 - v)f(\psi_-) + vf(\psi_+) - f(\phi) \\
=& (1 - v)f(\psi_-) + vf(\psi_-) + vk^\alpha(\partial_\alpha f)_{\psi_-} - f(\phi) \\
=& f(\psi_-) + vk^\alpha(\partial_\alpha f)_{\psi_-} - \sum_{i=0}^\infty \frac{1}{i!}\left[(vk^\alpha \partial_\alpha)^i f\right]|_{\psi_-} \\
=& -\frac{1}{2}(vk^\alpha)(vk^\beta)(\partial_\alpha \partial_\beta f)|_{\psi_-} + \mathcal{O}[(vk)^3].
\end{aligned} \tag{S.14}$$

The leading term in the stabilisation energy $\Delta f$ thus involves the Hessian $\partial_\alpha\partial_\beta f$. To this order $\Delta f$ is always negative given all eigenvalues of the Hessian $\partial_\alpha\partial_\beta f$ are positive, since the system is locally stable near the binodal boundary. Graphically, $\Delta f$ encodes the difference in free energy density between $f(\phi)$ and that arising from the common-tangent construction. In experimental settings the phase-separating system can typically be probed close to the dilute phase boundary so a second-order approximation is sufficient.

#### III. EXPRESSION FOR $n_\alpha$

We use the equilibrium equations Eqs. (S.12a) and (S.12b), applying perturbations to $\psi_-^\alpha$ and $\psi_+^\alpha$, we obtain

$$\delta\psi_-^\beta(\partial_\alpha\partial_\beta f)|_{\psi_-} = \delta\psi_+^\beta(\partial_\alpha\partial_\beta f)|_{\psi_+} \tag{S.15}$$

and

$$\psi_-^\alpha\delta\psi_-^\beta(\partial_\alpha\partial_\beta f)|_{\psi_-} = \psi_+^\alpha\delta\psi_+^\beta(\partial_\alpha\partial_\beta f)|_{\psi_+}. \tag{S.16}$$

Substituting Eq. (S.15) into Eq. (S.16) we get

$$\delta\psi_-^\beta k^\alpha(\partial_\alpha\partial_\beta f)|_{\psi_-} = 0 \tag{S.17}$$

so we can then make the identification

$$n_\beta \propto k^\alpha(\partial_\alpha\partial_\beta f)|_{\psi_-} \tag{S.18}$$

up to a normalisation constant.

#### IV. COMPUTATIONAL MODEL TO ILLUSTRATE $D^{(\alpha)}$ IN THE FULL PHASE SPACE

We use the Flory-Huggins free energy density [4, 5] with 2 solutes of unit length and implicit solvent:

$$f_{\rm FH} = \phi^1\ln\phi^1 + \phi^2\ln\phi^2 + \frac{1}{2}\chi_{11}\phi^1\phi^1 + \frac{1}{2}\chi_{22}\phi^2\phi^2 + \chi_{12}\phi^1\phi^2 + (1-\phi^1-\phi^2)\ln\big(1-\phi^1-\phi^2\big) \tag{S.19}$$

where the first two terms are solute entropies, the three terms following are homotypic interactions and the heterotypic interaction, and the last term is solvent entropy. The interaction energy values used to generate plots in the main text are summarised in Table I.

To compute the full phase diagram, we first generate a grid of 1000 by 1000 points with coordinates $(\phi^1, \phi^2)$ with $0 < \phi^{1,2} < 1$. By computing $f_{\rm FH}$ at each of these points and applying the convexhull algorithm to the 3-dimensional point set $(f_{\rm FH}, \phi^1, \phi^2)$ a set of simplexes are generated. Phase-separated regions are determined by comparing the projection of each simplex onto the $(\phi^1, \phi^2)$ space with the grid size, and the tie line can be determined from the long axis of the simplex [6].

After the dilute phase boundary and tie lines are calculated, we compute $D^1$ by first evaluating the ratio

$$\frac{D^1}{D^2} = \left(\frac{n_1}{n_2}\right)\left(\frac{k^1}{k^2}\right) \tag{S.20}$$

where we identify the first round bracket as the normal vector for the dilute phase boundary and the second round bracket as the tie line gradient. We next use $D^2 = 1 - D^1$ to obtain an expression for $D^1$:

$$D^1 = \frac{D^1/D^2}{1 + D^1/D^2}. \tag{S.21}$$

| Plot | $\chi_{11}$ | $\chi_{22}$ | $\chi_{12}$ |
|---|---|---|---|
| Fig. 2a | −4.3 | −4.3 | −3.9 |
| Fig. 2b | −4.3 | −4.3 | −4.3 |
| Fig. 2c | −4.3 | −4.3 | −4.9 |
| Fig. 2d | −5.2 | −4.2 | −5.8 |
| Fig. 2e | −4.2 | −5.2 | −5.8 |
| Fig. 3a | 0 | 0 | −10 |
| Fig. 3b | 0 | 0 | 8 |

TABLE I. Summary of parameters used to generate full phase diagrams.

#### V. RESPONSE FUNCTION DERIVATION

Starting from the mass conservation

$$\phi^\alpha = (1-v)\psi_-^\alpha + v\psi_+^\alpha \tag{S.22}$$

we first use the tie line definition $k^\alpha = \psi_+^\alpha - \psi_-^\alpha$ to write

$$\phi^\alpha = \psi_-^\alpha + vk^\alpha. \tag{S.23}$$

Applying a small perturbation, this is

$$\delta\phi^\alpha = \delta\psi_-^\alpha + k^\alpha\delta v + v\delta k^\alpha \tag{S.24}$$

where $\delta k^\alpha = \delta\psi_+^\alpha - \delta\psi_-^\alpha$. To make progress, we multiply both sides by $n_\alpha$ and contracting, noting $n_\alpha$ satisfies, by definition, $n_\alpha\delta\psi_-^\alpha = 0$. This gives an expression for $\delta v$:

$$\delta v = \frac{n_\alpha\delta\phi^\alpha - vn_\alpha\delta k^\alpha}{n_\gamma k^\gamma}. \tag{S.25}$$

Substituting $\delta v$ into Eq. (S.24) we then obtain

$$\delta\psi_-^\alpha + \left(\delta_\beta^\alpha - \frac{n_\beta k^\alpha}{n_\gamma k^\gamma}\right)\delta k^\beta v = \left(\delta_\beta^\alpha - \frac{n_\beta k^\alpha}{n_\gamma k^\gamma}\right)\delta\phi^\beta. \tag{S.26}$$

Eq. (S.26) quantifies how the dilute phase $\psi_-^\alpha$ responds to changes in overall concentration of each component $\phi^\beta$. Eq. (S.26) takes a simpler form in two special cases: 1) when $v = 0$, corresponding to probing the response right at the phase boundary, terms involving $\delta k^\alpha$ drops out; 2) $\delta k^\alpha$ is parallel to $k^\alpha$, in which case we can write $\delta k^\alpha = Ak^\alpha$ for some constant $A$, and it is easy to check $\left(\delta_\beta^\alpha - \frac{n_\beta k^\alpha}{n_\gamma k^\gamma}\right)k^\beta = 0$. In both of these cases we arrive at the simpler equation

$$\delta\psi_-^\alpha = \left(\delta_\beta^\alpha - \frac{n_\beta k^\alpha}{n_\gamma k^\gamma}\right)\delta\phi^\beta. \tag{S.27}$$

#### VI. EXPERIMENTAL PROCEDURE FOR HOMOTYPIC RESPONSE FUNCTION MEASUREMENT OF POLY(A) RNA

Poly(A) RNA with molecular weight of 700-3500 kilodaltons is purchased from Sigma, and a stock solution of 4g/l prepared by dissolving lyophilised powder in Milli-Q water. Polyethylene glycol (PEG) with molecular weight of 10 kilodaltons is purchased from Sigma and dissolved in 1M KCl, 50mM HEPES at pH 7.3 at 20% by weight. The working solutions are at 8% PEG, 1.6M or 1.7M KCl, 50mM HEPES, and pH 7.3. These are prepared by mixing the 20% PEG solution with solutions of 4M KCl, 50mM HEPES, and 0M KCl, 50mM HEPES.

Experimental samples are prepared by mixing the 8% PEG working solution, poly(A) stock solution and Milli-Q water in PCR tubes, with a total volume of 20µl. After mixing and vortexing, the samples are spun down at 13400 rpm for 2 minutes and 2µl of the supernatant is taken from each sample to measure the dilute phase [poly(A)] in a NanoDrop machine (ThermoFisher).


[1] J. W. Gibbs, On the equilibrium of heterogeneous substances, American Journal of Science and Arts **16**, 441 (1874).
